# Nisin G is a novel nisin variant produced by a gut-derived *Streptococcus salivarius*

**DOI:** 10.1101/2022.02.15.480493

**Authors:** Garreth W. Lawrence, Enriqueta Garcia-Gutierrez, Calum J. Walsh, Paula M. O’Connor, Máire Begley, Paul D. Cotter, Caitriona M. Guinane

## Abstract

*Fusobacterium nucleatum* is an emerging human pathogen associated with a number of intestinal conditions including colorectal cancer (CRC) development. Screening for gut-derived strains that exhibit anti-*F. nucleatum* activity revealed *Streptococcus salivarius* DPC6487 as a strain of interest. Whole genome sequencing analysis of DPC6487 resulted in the identification of a gene predicted to encode a novel nisin variant designated nisin G. The structural nisin G peptide differs from the prototypical nisin A with respect to seven amino acids (Ile4Tyr, Ala15Val, Gly18Ala, Asn20His, Met21Leu, His27Asn and His31Ile), including differences that have not previously been associated with a natural nisin variant. The nisin G gene cluster consists of *nsgGEFABTCPRK* with transposases encoded between the nisin G structural gene (*nsgA*) and *nsgF*. The cluster lacked an equivalent to the *nisI* immunity determinant. *S. salivarius* DPC6487 exhibited a narrower spectrum of activity compared to the nisin A producer, *Lactococcus lactis* NZ9700, when assessed through deferred antagonism-based assays. Such narrow spectrum activity is desirable as it is less likely to lead to collateral damage to gut commensals.

Ultimately, this is the first report of a nisin variant produced by a representative of a species that is frequently a focus for probiotic development. The production of this bacteriocin by a gut-derived *S. salivarius* and its narrow spectrum activity against *F. nucleatum* indicates that this strain merits further attention to determine its potential for probiotic-related applications.

## Introduction

The human gut microbiome comprises trillions of microbes, some members of which coexist while others compete for essential resources that determine their survival^1^. Co-evolution and competition have resulted in the development of several mechanisms that aid in the survival of microorganisms, including the secretion of antimicrobial peptides^2–4^. For this reason, it is not surprising that the gut microbiome is regarded as a reservoir of antimicrobial peptides such as bacteriocins that hold therapeutic potential^5–7^. Bacteriocins are ribosomally-synthesized antimicrobial peptides that display a narrow- or broad-spectrum of activity^8^. As antibiotic resistant pathogens continue to emerge, bacteriocins have increasingly been studied as potential antimicrobial alternatives^9^. Furthermore, bacteriocin-producing probiotic bacteria have the potential to target disease-associated taxa *in situ* in the gut^10^.

The genus *Streptococcus* is well-known for its bacteriocin-producing potential, including the production of lantibiotics^11^. The prototypical lantibiotic is nisin. First discovered in *Lactococcus lactis* in 1928, variants of the broad spectrum lantibiotic nisin A have since been reported to be produced by *Streptococcus* species, i.e., strains of *Streptococcus uberis* produces nisin variants nisin U and U2, nisin H is produced by a strain of *Streptococcus hyointestinalis*^12^, while nisin P is produced by a strain of *Streptococcus agalactiae*^5^. Interestingly, a nisin-like peptide, salivaricin D, is produced by a strain of *Streptococcus salivarius*. However, salivaricin D differs greatly from nisin A^13^. Nisin exerts its antimicrobial activity through pore formation and inhibition of cell wall biosynthesis^14^. Nisin A was approved as a food preservative in 1953^15^ and in 1988 the FDA granted nisin generally regarded as safe (GRAS) status. Furthermore, nisin has been investigated to assess its biotherapeutic potential, including the targeting of bacterial pathogens associated with cancer^16^.

Several strains of *S. salivarius* produce the lantibiotic salivaricin A, with five variants having been identified to date^17^. Notably, *S. salivarius* strain K12, a salivaricin B and salivaricin A2 co-producer with antagonistic activity against the pathogen *Streptococcus pyogenes*, has been developed as a commercial probiotic and has passed rigorous safety assessment for human use^18,19^. As strains of *S. salivarius* have been shown to benefit human health in numerous clinical trials^20–22^, there is considerable merit in screening other strains for traits with a view to the further development of novel probiotics with antimicrobial activity^23^.

*Fusobacterium nucleatum* is an emerging human pathogen shown to be associated with colorectal cancer (CRC), with high abundances of *Fusobacterium* and *F. nucleatum* having been identified at higher relative abundance in the faecal, tissue and mucosal samples of patients with CRC, relative to the corresponding samples from healthy controls^24–30^. Furthermore, increased abundances of *F. nucleatum* has been found to correlate with poor patient prognosis^31^. Indeed, evidence suggests that *F. nucleatum* contributes to the development of CRC^32^, and thus represents a potential therapeutic target^10^

In this study, an intestinal isolate of *S. salivarius* DPC6487, previously isolated from a neonatal faecal sample^6^, was found to demonstrate activity against strains of *F. nucleatum*. Further analysis suggests this strain exhibits a narrow spectrum of antimicrobial activity. Genome sequencing revealed a gene cluster encoding a novel nisin variant, designated nisin G. This gene cluster was analysed, and it, and the antimicrobial spectrum of *S. salivarius* DPC6487, were compared with those of a nisin A producer.

## Results

### Characterisation of an unknown antimicrobial produced by S. salivarius DPC6487

*S. salivarius* DPC6487 was previously isolated from a neonatal faecal sample in a screening study of the human gut microbiota and was previously reported to demonstrate antimicrobial activity against a strain of *Lactobacillus bulgaricus*^6^. In this study, further analysis of *S. salivarius* DPC6487 revealed that it demonstrates antimicrobial activity against the pathogen *F. nucleatum* DSM15643 **(Figure 1A)**. MALDI-TOF colony mass spectrometry of *S. salivarius* DPC6487 revealed the presence of a molecule with a mass of 3405 Da **(Figure 1B)**.

**Figure 1.**
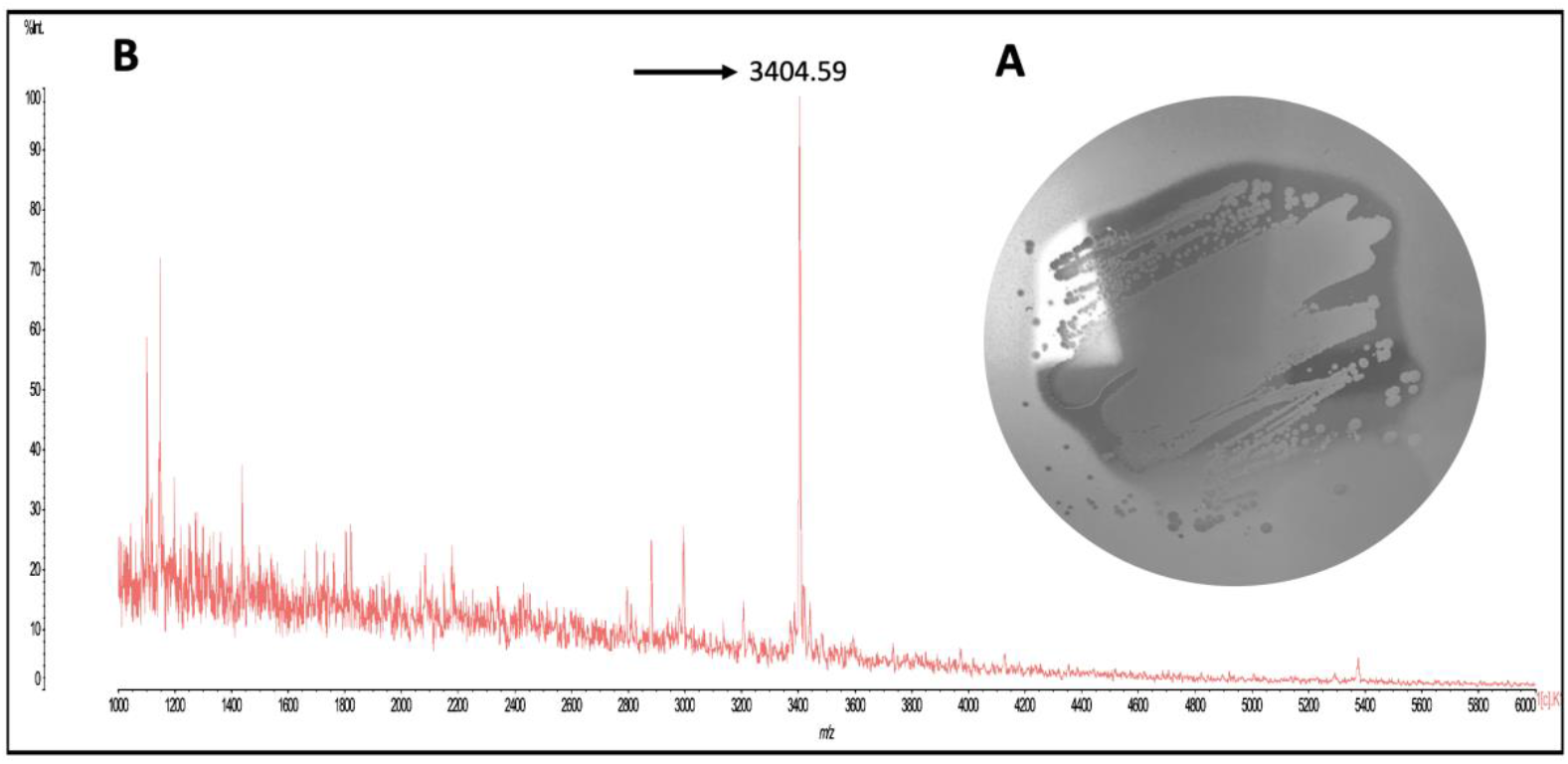
An unknown antimicrobial is produced by *S. salivarius* DPC6487. Deferred antagonism assay whereby *S. salivarius* DPC6487 demonstrated antimicrobial activity against *F. nucleatum* DSM15643 **(A)**. The presence of a 3404.59 Da mass was revealed by MALDI-TOF MS **(B)**.

The antimicrobial activity of cell free supernatant (CFS) from an overnight culture of *S. salivarius* DPC6487 remained stable after treatment at 37, 60, 70, 80, 90, 100, and 121°C for 10 minutes **(Figure 2A)**. The activity of the CFS also remained active at pH 2, 3, 5, 8, and 10, with antimicrobial activity being greatest at pH 2-3, and decreasing from pH 5 to pH 10 **(Figure 2B)**. Proteinase treatment of *S. salivarius* DPC6487 CFS resulted in the loss of antimicrobial activity as previously reported^6^ and, taken together, the results indicate that the antimicrobial being produced was heat stable, proteinaceous in nature, and retained activity in acidic environments **(Figure 2C)**.

**Figure 2.**
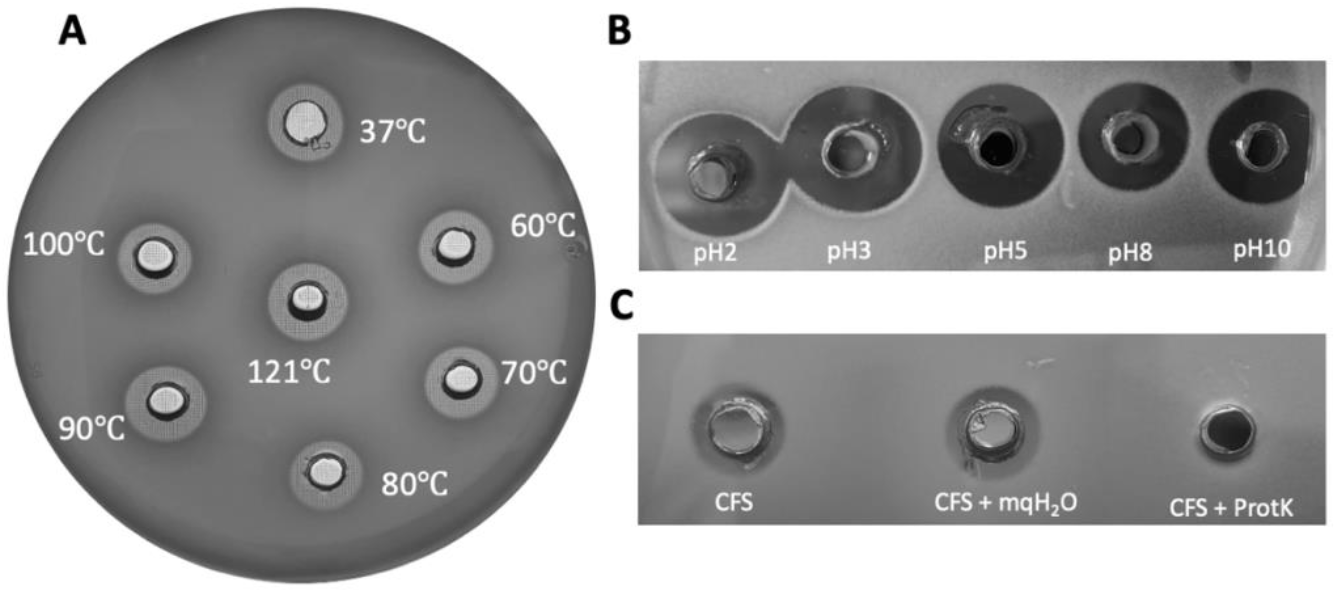
Effect of heat (A), pH (B) and proteinase K (C) on the CFS of *S. salivarius* DPC6487. After subjection to heat, pH and proteinase K the antimicrobial activity of *S. salivarius* DPC6487 CFS was assessed against *L. delbrueckii* ssp. *bulgaricus* DPC5383.

### Whole genome sequencing of S. salivarius DPC6487 revealed a gene cluster encoding a natural nisin variant

Sequencing of *S. salivarius* DPC6487 yielded a 2,240,822 bp draft genome with an overall GC content of 39.71%. Blast analysis of the 16S rRNA gene confirmed a 99% identity to *S. salivarius* 16S rRNA gene sequences. Contig analysis via the bacteriocin mining tool BAGEL4 indicated the presence of a potential nisin variant. Sequencing analysis confirmed that the genome of *S. salivarius* DPC6487 harboured a gene predicted to encode a natural nisin variant that was designated nisin G. The putative novel nisin G peptide contains the following amino acid substitutions relative to nisin A: Ile4Tyr, Ala15Val, Gly18Ala, Asn20His, Met21Leu, His27Asn and His31Ile **(Figure 3)**. Nisin G is most similar to nisin Q (produced by *L. lactis* 61-14) and share three amino acid substitutions when compared to nisin A: Ala15Val, Meth21Leu, and His27Asn, while differing with respect to five amino acids: Ile4Try, Gly18Ala, Asn20His, Val30Ile and His31Ile. Interestingly, nisin H, produced by *S. hyointestinalis*, shares no common amino acid substitutions with nisin G when compared to nisin A. However, amino acids at positions 18, 21 and 31 are changed in both variants. Nisin G contains three unique amino acids when compared to all natural nisin variants, i.e., Gly18Ala, Asp20His and His31Ile **(Figure 3)**.

**Figure 3.**
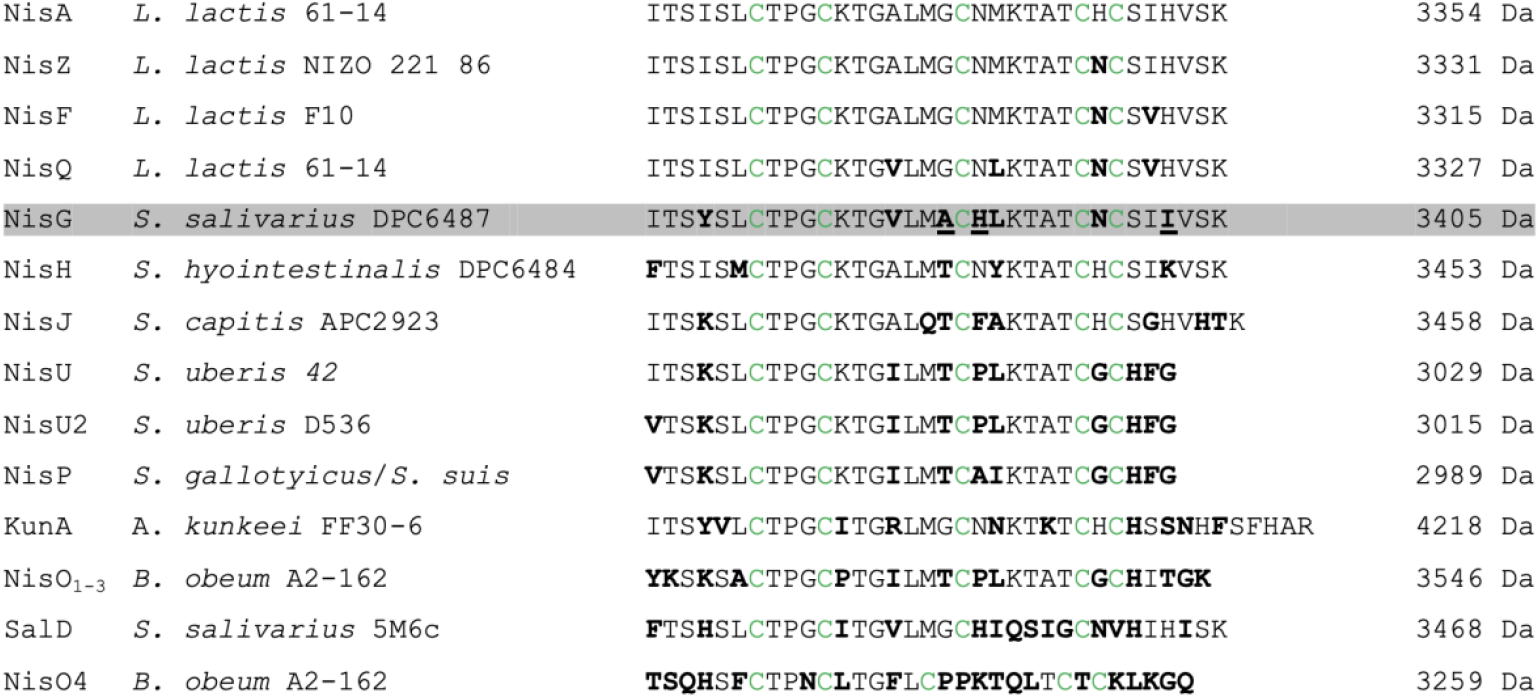
Sequence alignment of natural nisin (Nis) variants. Amino acid substitutions are highlighted in bold; cysteines are highlighted in green and nisin G (NisG) is shaded. Amino acid substitutions unique to NisG are underlined. Salivaricin D (SalD) produced by *S. salivarius* 5M6c and Kunkecin A (KunA) produced by *Apilactobacillus kunkeei* FF30-6 were included as they are considered ‘nisin-like’. Nisin accession (reference is provided where accession is not linked to primary source): Nisin A, ABN45880; Nisin Z, ABV64387; Nisin F, ABU45463; Nisin Q, ADB43136; Nisin H, AKB95119; Nisin J,^45^; Nisin U, Q2QBT0; Nisin U2, ABO32538; Nisin P,^58^; Nisin O,^34^; Kunkecin A^59^, Salivaricin D, AEX55166.

A phylogenetic comparison of all nisin variants demonstrated that respective streptococcal and lactococcal nisin variants group together. The *Blautia*-derived nisin O_1-4_ and salivaricin D are more distantly related **(Figure 4)**.

**Figure 4.**
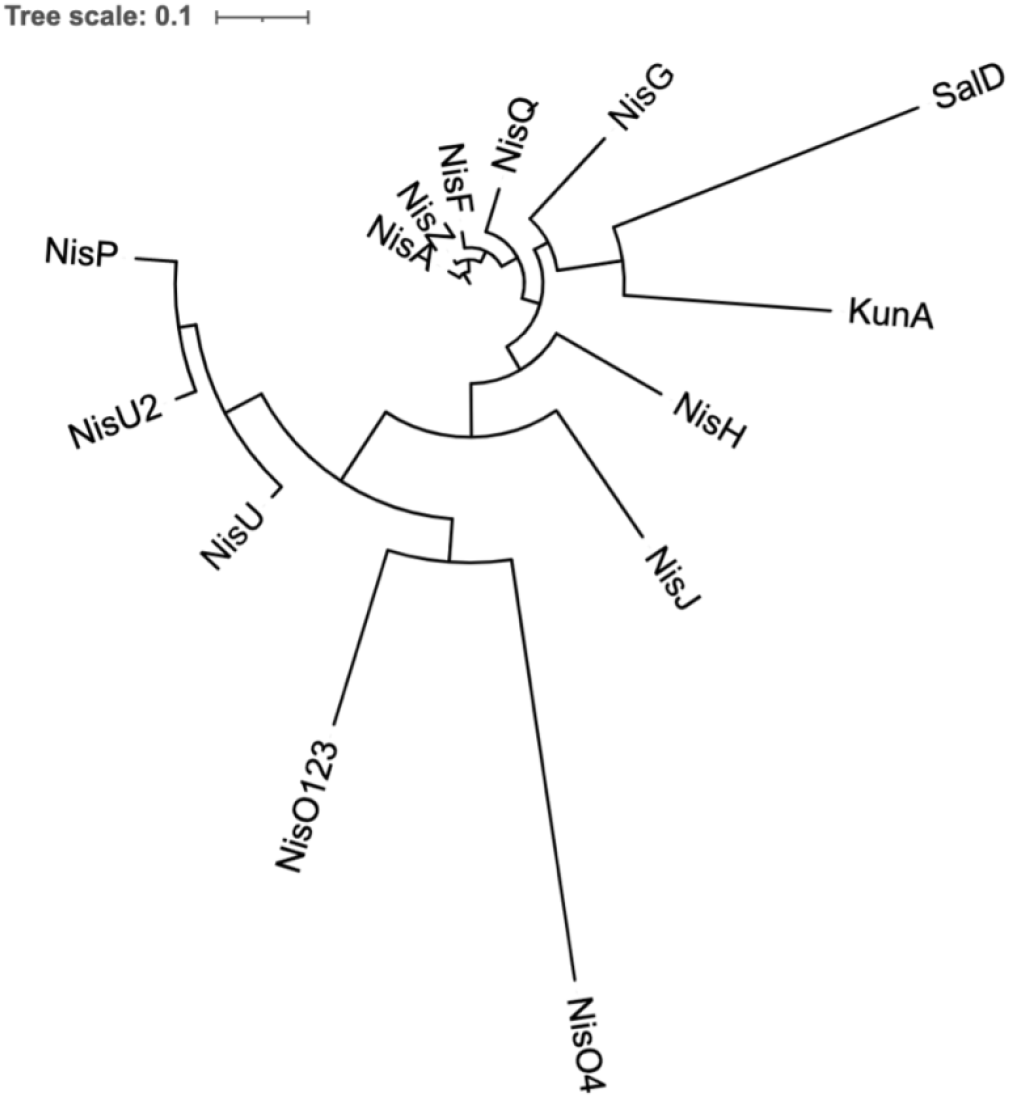
Phylogenetic relationship of nisin G to other natural nisin variants. The phylogenetic relationship was inferred using the neighbourhood-joining method^55^; evolutionary distances were computed using the Poisson correction method. All steps involved in the construction of the tree were performed in MEGAX^56^ and visualised using ITOL^57^. Branch length (Scale = 0.1) indicates the number of amino acid substitutions per site. Nis, Nisin; Sal, salivaricin; Kun, Kunkecin.

### Genomic analysis of S. salivarius DPC6487 identified genes putatively responsible for production of and immunity to nisin G

Genomic analysis identified nisin-related genes on a single assembled contig from the draft genome of *S. salivarius* DPC6487. The nisin G encoding gene (designated as *nsgA*) and associated biosynthesis and immunity genes were identified in a gene cluster spanning a region of 14.7 kb. The putative immunity determinants displayed a similar organisation to other nisin gene clusters including nisin U^33^, nisin O^34^ and nisin P^5^ gene clusters **(Figure 5)**. However, a notable feature of the nisin G genetic organisation is that *nsgFEG* are transcribed in the opposite direction to *nsgA*. Streptococcal transposases were found between the nisin G encoding gene *nsgA* and *nsgF*. A region encoding transposases was also evident outside the nisin G gene cluster adjacent to the immunity determinant *nsgG* (data not shown). The presence of transposases are also common in the genetic organisation with the cluster encoding the *S. hyointestinalis-*associated nisin, where a transposase is located between *nshP* and *nshR* and between *nshK* and *nshG*^12^ **(Figure 5)**. A transposase is also found between *slvG* and *slvK* and adjacent to *slvD* in the salivaricin D genetic organisation^13^. Similar to nisin H, the nisin G gene cluster lacks an equivalent to the immunity gene *nisI* and blast analysis against the entire *S. salivarius* DPC6487 genome confirmed the absence of a *nisI* homologue. Therefore, the organisation of nisin G gene cluster is *nsgGEFABTCPRK* **(Figure 5)**. The predicted nisin G pre-peptide sequence is composed of 57 amino acids containing a leader sequence of 23 amino acids. The nisin G mature peptide displays 78.95% and 84.21% amino acid identity to nisin A and nisin H, respectively, and only 70.18% to salivaricin D. Furthermore, the nisin G biosynthesis and immunity genes demonstrated highest amino acid identity to the products of the equivalent nisin H genes, ranging from 89% to 98% identity **(Figure 5)**.

**Figure 5.**
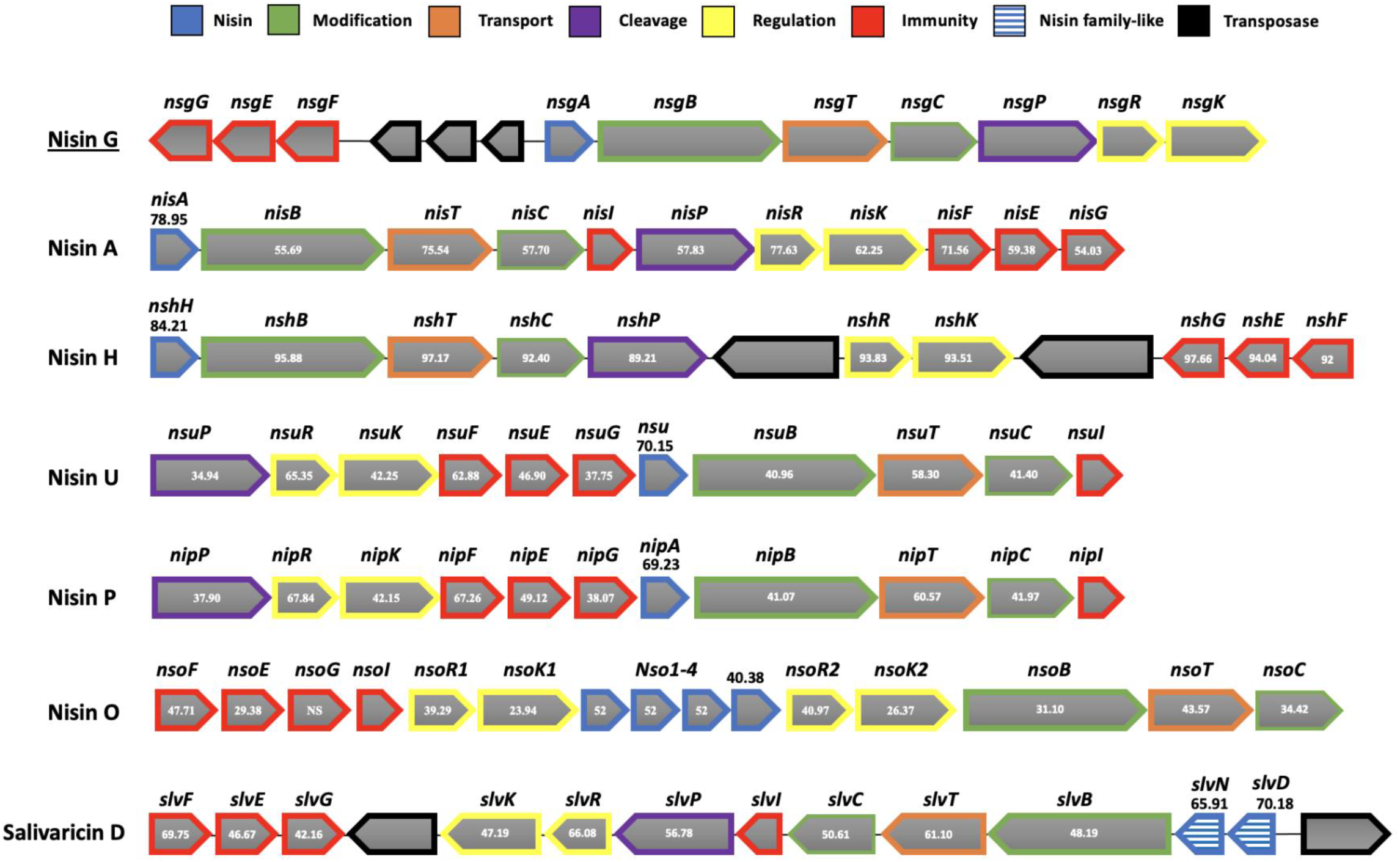
Comparison of the nisin G gene cluster found in the genome of *S. salivarius* DPC6487 with nisin A, nisin H, nisin U, nisin P, nisin O and salivaricin D gene clusters. Figures shown in nisin A, nisin H and salivaricin D gene clusters represent percentage amino-acid identity to nisin G gene homologues. Where no percentage identity is indicated, no homologue was identified in the nisin G cluster. NS, no similarity.

### Spectrum of inhibition of S. salivarius DPC6487 and L. lactis NZ9700

The spectrum of inhibition of nisin G-producing *S. salivarius* DPC6487 and nisin A-producing *L. lactis* NZ9700 was evaluated by deferred antagonism assay **(Table 2)**. *L. lactis* NZ9700 demonstrated bioactivity (i.e., representing the combined impact of differences in production levels and specific activity) against all *Streptococcus* species, however, of the streptococci tested, *S. salivarius* DPC6487 was only active against *S. thermophilus* strains DPC5473 and DPC5657 and *S. uberis* DPC4344. Notably, *S. salivarius* DPC6487 showed increased bioactivity against *S. agalactiae* ATCC13813. In contrast to *L. lactis* NZ9700, *S. salivarius* DPC6487 showed no bioactivity against *S. mutans* strains DPC6160 and DPC6161 or *S. simulans* APC3482. *S. salivarius* DPC6487 did not demonstrate activity against the *Lactobacillus, Listeria*, or *Staphylococcus* strains tested here, which contrasted with the distinct antimicrobial activity observed when *L. lactis* NZ9700 was used to target these genera. As nisin G-producing *S. salivarius* DPC6487 originally showed antimicrobial activity against *F. nucleatum* DSM15643, it was hypothesised that activity would be observed against other *Fusobacterium* species and *F. nucleatum* strains. When tested, it was established that *S. salivarius* DPC6487 and *L. lactis* NZ9700 both showed antimicrobial activity against *F. nucleatum* strains DSM15643, DSM19508, and DSM19507. *Fusobacterium periodonticum* DSM19545 was also susceptible to *S. salivarius* DPC6487 and *L. lactis* NZ9700, however, no activity was observed against *E. coli* K12 or *E. coli* ATCC25927. It was also established that *L. lactis* NZ9700, but not *S. salivarius* DPC6487, demonstrated bioactivity against *Clostridioides difficile* DPC6357 **(Table 2)**.

## Discussion

The use of bacteriocin-producing probiotics to target disease-associated taxa as an alternative to antibiotics is gaining ever more interest^35^. Furthermore, specific species such as *S. salivarius* represent promising candidates for probiotic development with a view to targeting pathogenic bacteria^23^.

In this study, a novel nisin variant, designated nisin G, produced by a gut isolate of *S. salivarius*, was identified. The producing strain, DPC6487, was shown to have antimicrobial activity against the pathogen *F. nucleatum*. Colony mass spectrometry of *S. salivarius* DPC6487 revealed a mass of 3405 Da, which indicated the potential secretion of a bacteriocin. Additionally, heat, pH and proteinase sensitivity assays with *S. salivarius* DPC6487 CFS indicated that the antimicrobial being produced was likely proteinaceous. Indeed, many bacteriocins such as lantibiotics are heat-stable and remain active in highly acid environments^36,37^.

Genome sequencing of *S. salivarius* DPC6487 revealed an operon with a gene encoding a novel nisin variant, designated nisin G. The nisin variant nisin G contains seven amino acid substitutions compared to nisin A^38^ and has three amino acid substitutions that are distinct from all other reported natural nisin variants; an alanine at position 18, a histidine at position 20 and an isoleucine at position 31. Notably, this is the first nisin reported from the species *S. salivarius*. However, streptococcal-derived nisin variants are common. For example, *S. agalactiae, S. uberis, and S. hyointestinalis* produce the variants nisin P^5^, nisin U^33^, and nisin H ^12^, respectively. Interestingly, a nisin-like peptide, salivaricin D, has also been reported in *S. salivarius* 5M6c^13^. However, salivaricin D differs greatly from nisin A, i.e., by 14 amino acids, contains a unique region of 6 adjacent amino acids (HIQSIG) and comprises only 4 ring structures, lacking the region referred to as ring E that is found in nisin molecules.

CRC-associated *F. nucleatum* represents a potential therapeutic target, and eradicating or suppressing the growth of this pathogen within the human gut microbiome may ultimately contribute to reducing or removing the overall risk of disease development^10^. Both nisin producers *S. salivarius* DPC6487 and *L. lactis* NZ9700 displayed distinct bioactivity against *F. nucleatum* DSM15643. A limited number of studies have evaluated the antimicrobial activity of nisin against oral pathogens, which included *F. nucleatum, in vitro*^39,40^.

Overall, *S. salivarius* DPC6487 showed a narrower spectrum of activity compared to *L. lactis* NZ9700, using the overlay method, with activity against just *Fusobacterium spp*. and other streptococci. Furthermore, *S. salivarius* DPC6487 showed no activity against *C. difficile* or *E. coli* strains tested in this study, indicating a potential narrow-spectrum of activity. Narrow-spectrum antimicrobials are of particular interest as alternatives to antibiotics as they leave the surrounding microbiota unharmed^41^. Both *S. salivarius* DPC6487 and *L. lactis* NZ9700 inhibited two additional *F. nucleatum* strains DSM19508 and DSM19509 and a *F. periodonticum* strain DSM19545, further supporting the anti-*Fusobacterium* potential of nisin and nisin-producers. In addition, it was noted that *S. agalactiae* ATCC13813 was most susceptible of all streptococci tested to *S. salivarius* DPC6487. As invasive infections caused by *S. agalactiae* have been reported in pregnant women, new-borns and in adults with immunosuppressive diseases^42^, this finding suggests a possible application for *S. salivarius* DPC6487 to reduce the risk of *S. agalactiae*–associated illnesses. As expected, the nisin A-producing *L. lactis* NZ9700 showed a broad spectrum of activity with activity against *Fusobacterium, Clostridioides, Streptococcus, Lactobacillus, Staphylococcus*, and *Listeria*. Furthermore, no activity was observed against Gram negative *E. coli* strains, as previously reported for nisin variants^5,12^. The narrower-spectrum of activity observed for *S. salivarius* DPC6487 may be a consequence of the unique amino acid substitutions present in nisin G, however, further analysis is required to support this. Indeed, site directed mutagenesis of the hinge region of nisin A resulting in either histidine at position 20 or leucine at position 21 (which are observed naturally in nisin G) reduced the bioactivity of the peptide against Gram-positive pathogens^43^. Therefore, histidine in position 20 together with leucine in position 21 of the nisin G molecule may be contributing to the decreased potency observed compared to nisin A. Further analysis is required to fully understand the effect of these amino acid substitutions on the activity of nisin G.

In conclusion, numerous studies have used nisin as a means of targeting disease-associated taxa^16^. As such, nisin-producing bacteria represent potential candidates for the development of antimicrobial-producing probiotics, which may be utilized to target pathogenic bacteria, and consequently, lower the overall risk of disease development. The nisin G producing *S. salivarius* DPC6487 is a candidate for probiotic development. *In vivo* trials and safety assessments are needed to further elucidate the potential of *S. salivarius* DPC6487 as a potential probiotic.

## Materials and Methods

### Bacterial strains and cultivation media

*S. salivarius* DPC6487, previously isolated from a neonatal faecal sample, was sourced from the Teagasc culture collection^6^. The strain was cultivated under anaerobic conditions at 37°C in Brain Heart Infusion (BHI, Difco Laboratories, Detroit, MI, USA) broth and agar medium containing 1.5% w/v agar. Anaerobic conditions were simulated using anaerobic jars with Anaerocult A gas packs (Merck, Darmstadt, Germany) or a Don Whitley Anaerobic workstation (nitrogen 85%, carbon dioxide 5%, hydrogen 10%). A full list of bacteria and their culture conditions used in this study are presented in **Table 1**.

**Table 1.** Bacterial strains and their culture conditions used in this study

| Indicator organism | Culture Medium | Temp (°C) + Conditions |
| --- | --- | --- |
| *Fusobacterium nucleatum* DSM15643 | FAA | 37, Anaerobic |
| *Fusobacterium nucleatum* DSM19507 | FAA | 37, Anaerobic |
| *Fusobacterium nucleatum* DSM19508 | FAA | 37, Anaerobic |
| *Fusobacterium periodonticum* DSM19545 | FAA | 37, Anaerobic |
| *Escherichia coli* K12 | LB | 37, Aerobic |
| *Escherichia coli* ATCC25927 | LB | 37, Aerobic |
| *Clostridioides difficile* DPC6357 | FAA | 37, Anaerobic |
| *Streptococcus agalactiae* DPC7040 | BHI | 37, Anaerobic |
| *Streptococcus uberis* DPC4344 | BHI | 37, Anaerobic |
| *Streptococcus mutans* DPC6160 | BHI | 37, Anaerobic |
| *Streptococcus mutans* DPC6161 | BHI | 37, Anaerobic |
| *Streptococcus thermophilus* DPC5472 | BHI | 37, Anaerobic |
| *Streptococcus thermophilus* DPC5657 | BHI | 37, Anaerobic |
| *Streptococcus agalactiae* ATCC13813 | BHI | 37, Anaerobic |
| *Streptococcus simulans* APC3482 | BHI | 37, Anaerobic |
| *Listeria monocytogenes* DPC3564B | BHI | 37, Aerobic |
| *Listeria monocytogenes* DPC3853B | BHI | 37, Aerobic |
| *Listeria innocua* DPC3572 | BHI | 37, Aerobic |
| *Lactobacillus fermentum* DPC3320 | BHI | 37, Aerobic |
| *Lactobacillus plantarum* DPC6667 | BHI | 37, Aerobic |
| *Lactobacillus delbrueckii ssp. bulgaricus* DPC5383 | MRS | 37, Anaerobic |
| *Lactococcus lactis* NZ9700 | GM17 | 30, Aerobic |
| *Staphylococcus aureus* DPC7016 | BHI | 37, Aerobic |
| Methicillin-resistant *Staphylococcus aureus* DPC5654 | BHI | 37, Aerobic |
| *Staphylococcus epidermidis* DPC5990 | BHI | 37, Aerobic |

FAA, Fastidious Anaerobic Agar (Lab M, Lancashire, UK); BHI, Brain Heart Infusion; GM17, Glucose (0.5%) M17; (Difco Laboratories, Detroit, MI); LB, Luria-Bertani Medium; MRS, de Man, Rogosa, and Sharpe medium (Difco Laboratories, Detroit, MI); DSMZ, German Collection of Microorganisms and Cell Culture GmbH; ATCC, American Type Culture Collection; APC, Alimentary Pharmabiotic Centre; DPC, Teagasc Culture Collection.

**Table 2.** Indicators used in the inhibitory activity spectrum assessment of *S. salivarius* DPC6487 and *L. lactis* NZ9700 and the degree of inhibition. Zones of inhibition (mm) were measured around single colonies and relative sensitivity determined. +: zones of size <2mm, ++: zones of size 2-5mm, +++: zones of size >5mm.

| Indicator | Inhibition |  |
| --- | --- | --- |
|  | *S. salivarius*<br>DPC6487<br>(nisin G producer) | *L. lactis*<br>NZ9700<br>(nisin A producer) |
| *Fusobacterium nucleatum* DSM15643 | ++ | ++ |
| *Fusobacterium nucleatum* DSM19507 | ++ | ++ |
| *Fusobacterium nucleatum* DSM19508 | ++ | ++ |
| *Fusobacterium periodonticum* DSM19545 | ++ | ++ |
| *Escherichia coli* K12 | - | - |
| *Escherichia coli* ATCC25927 | - | - |
| *Clostridioides difficile* DPC6357 | - | + |
| *Streptococcus mutans* DPC6160 | - | + |
| *Streptococcus mutans* DPC6161 | - | + |
| *Streptococcus thermophilus* DPC5472 | + | +++ |
| *Streptococcus thermophilus* DPC5657 | + | +++ |
| *Streptococcus agalactiae* ATCC13813 | +++ | +++ |
| *Streptococcus uberis* DPC4344 | + | + |
| *Streptococcus simulans* APC3482 | - | +++ |
| *Lactobacillus delbrueckii* subsp. *bulgaricus* DPC5383 | +++ | +++ |
| *Lactobacillus fermentum* DPC3320 | - | + |
| *Lactobacillus plantarum* DPC6667 | - | +++ |
| *Staphylococcus aureus* DPC7016 | - | ++ |
| Methicillin-resistant *Staphylococcus aureus* DPC5654 | - | +++ |
| *Staphylococcus epidermidis* DPC5990 | - | +++ |
| *Listeria monocytogenes* DPC3564B | - | +++ |
| *Listeria monocytogenes* DPC3853B | - | ++ |
| *Listeria innocua* DPC3572 | - | + |

DPC, Teagasc Culture collection; APC, APC Culture Collection; DMS, German Collection of Microorganisms; ATCC, American Type Culture Collection.

### Colony mass spectrometry of S. salivarius DPC6487

Fully grown colonies of *S. salivarius* DPC6487 were mixed with 50μl 2-propanol 0.1% TFA, vortexed three times and centrifuged at 21,300 g for 30 seconds. MALDI TOF mass spectrometry was performed on the cell free supernatant using an Axima TOF^2^ MALDI-TOF mass spectrometer (Shimadzu Biotech, Manchester, UK). A 0.5μl aliquot of matrix solution (α-cyano 4-hydroxy cinnamic acid, 10 mg/ml in acetonitrile - 0.1% (v/v) TFA) was deposited onto the target and left for 5 seconds before being removed. The residual solution was allowed to dry and 0.5μl sample solution was deposited onto the pre-coated sample spot. A 0.5μl aliquot of matrix solution was added to the deposited sample and allowed to dry. The sample was subsequently analysed in positive-ion linear mode.

### Antimicrobial activity assays

Antimicrobial activity of *S. salivarius* DPC6487 and *L. lactis* NZ9700 against *F. nucleatum* DSM15643 and a range of indicator strains was determined by a deferred antagonism assay^44^. In brief, a fully cultured nisin-producer streak plate was overlayed with 0.75% agar seeded with the indicator microorganism and incubated according to the growth condition of the indicator microorganism **(Table 1)**. Inhibitory activity of *S. salivarius* DPC6487 cell-free supernatant (CFS) against *L. bulgaricus* DPC5383 was determined by well diffusion assay (WDA) as previously described^45^. Briefly, 50 μl of CFS prepared from 10 mL of an overnight culture of *S. salivarius* DPC6487 were added to wells of MRS agar plates seeded with *L. bulgaricus* DPC5383 and incubated at 37°C under anaerobic conditions for 18-20 hours. Antimicrobial activity was determined by the presence of a zone of inhibition.

### Heat, pH and protease sensitivity assays

The stability of the antimicrobial secreted by *S. salivarius* DPC6487 to a variety of physio-chemical factors was examined. Initially, CFS was subjected to temperatures of 37, 60, 70, 80, 90, 100 and 121°C for 10 minutes and the heat treated CFS was evaluated for its antimicrobial activity against *L. bulgaricus* DPC5383 by WDA, as previously described. A pH stability test was performed by altering the pH of *S. salivarius* DPC6487 CFS to 2.0, 3.0, 5.0, 8.0, and 10.0 by addition of 1M HCL or 1M NaOH. Antimicrobial activity of the pH adjusted CFS against *L. bulgaricus* DPC5383 was evaluated by WDA. A proteinase sensitivity assay was performed by incubating the *S. salivarius* DPC6487 CFS with 20 mg/mL proteinase K (Sigma) at 37°C for 1 hour. CFS mixed with an equal volume of molecular biology grade water was included as a control. The antimicrobial activity of these CFSs were determined against *L. bulgaricus* DPC5383 by WDA.

### S. salivarius DPC6487 whole genome sequencing and analysis

Genomic DNA was extracted from *S. salivarius* DPC6487 culture cell pellets using a GenElute™ Bacterial Genomic DNA Kit (Sigma-Aldrich; Co. Wicklow, Ireland). The purity and concentration of genomic DNA was confirmed using the NanoDrop 1000 (ThermoFisher Scientific, Dublin, Ireland) and Qubit® 2.0 Fluorometer (ThermoFisher Scientific, Dublin, Ireland) according to the respective protocols. DNA was prepared according to the Nextera XT DNA library preparation guide from Illumina and sequenced on an Illumina MiSeq (Teagasc, Moorepark Sequencing Facility). Quality trimming of the resulting raw FastQ files was performed using TrimGalore (v.0.6.0) (URL: https://www.bioinformatics.babraham.ac.uk/projects/trim_galore/), a wrapper script for cutadapt (v. 2.6)^46^ and FastQC (v. 0.11.8)^47^ with a q-score cut-off of 30 and minimum length after trimming of 105bp. Error correction and assembly into contigs was performed using SPAdes (v. 3.14)^48^ in “isolate” run mode. The assembled contigs of the draft genome were annotated using Prokka (v. 1.14)^49^ with RNAmmer^50^ for rRNA prediction. BAGEL4 software, an automated bacteriocin mining tool was used to detect the presence of putative bacteriocin operons^51^. Manual analysis of the contigs was then subsequently performed using the ARTEMIS genome browser ^52^. The putative bacteriocin gene clusters were manually annotated following sequence similarity analyses using the BLASTp algorithm and the non-redundant database provided by the NCBI ^53^ (http://blast.ncbi.nlm.nih.gov). Multiple sequence alignment of nisin amino acid sequences was performed using ClustalW^54^ and the phylogenetic relationship was inferred using the Neighbourhood Joining method^55^; evolutionary distances were computed using Poisson correction method and the tree was constructed with MEGAX with 1000 rounds of bootstrapping^56^ and visualised using ITOL^57^.

## Author Contributions

GWL wrote the main manuscript. CMG and PDC conceived the study idea and design. GWL performed the experiments and data analysis. CJW and EGG ran the bioinformatic pipeline. PMOC performed colony mass spectrometry. CMG, MB and PDC supervised, edited, and proofread the manuscript. All authors read and approved the final manuscript.

## Acknowledgements

The authors thank Harsh Mathur for excellent guidance, and Fiona Crispie and Laura Finnegan for their contributions to DNA sequencing.

## Data availability

The sequence of the nisin G gene cluster of *Streptococcus salivarius* DPC6487 will be deposited to GenBank

